## Supplementary Materials for "Approximate Bayesian computation supports a high incidence of chromosomal mosaicism in blastocyst-stage human embryos"

| | Dispersal = 0 | | Dispersal = 0.5 | | Dispersal = 1 | |
|---|---|---|---|---|---|---|
| | Meiotic | Mitotic | Meiotic | Mitotic | Meiotic | Mitotic |
| **Mean** | 0.40 | 0.063 | 0.57 | 0.021 | 0.58 | 0.015 |
| **Pctl. 2.5** | 0.34 | 0.054 | 0.54 | 0.018 | 0.55 | 0.014 |
| **Pctl. 25** | 0.38 | 0.059 | 0.56 | 0.020 | 0.57 | 0.015 |
| **Pctl. 50** | 0.40 | 0.062 | 0.57 | 0.020 | 0.58 | 0.015 |
| **Pctl. 75** | 0.42 | 0.065 | 0.58 | 0.021 | 0.59 | 0.016 |
| **Pctl. 97.5** | 0.45 | 0.071 | 0.60 | 0.023 | 0.60 | 0.017 |
| **MAP** | 0.40 | 0.063 | 0.57 | 0.020 | 0.58 | 0.015 |

**Supplementary Table 1: Inferred rates of meiotic and mitotic error at varying levels of dispersal.** The reported statistics are summaries of the corresponding posterior distributions. The 2.5 and 97.5 percentiles of the posterior distributions form the boundaries of the 95% credible intervals. The last row reports the maximum *a posteriori* (MAP) estimate (Capalbo et al. 2021).

| | Dispersal = 0 | | | Dispersal = 0.5 | | | Dispersal = 1 | | |
|---|---|---|---|---|---|---|---|---|---|
| | **Euploid** | **Mosaic** | **Aneuploid** | **Euploid** | **Mosaic** | **Aneuploid** | **Euploid** | **Mosaic** | **Aneuploid** |
| **Mean** | 0.23 | 0.19 | 0.58 | 0.23 | 0.19 | 0.58 | 0.23 | 0.19 | 0.58 |
| **Std. Dev.** | 0.0043 | 0.0031 | 0.0047 | 0.0032 | 0.0054 | 0.0056 | 0.0027 | 0.0048 | 0.0049 |
| **Min.** | 0.22 | 0.18 | 0.57 | 0.23 | 0.18 | 0.57 | 0.23 | 0.18 | 0.57 |
| **Pctl. 25** | 0.23 | 0.18 | 0.58 | 0.23 | 0.18 | 0.58 | 0.23 | 0.18 | 0.58 |
| **Pctl. 50** | 0.23 | 0.19 | 0.58 | 0.23 | 0.19 | 0.58 | 0.23 | 0.19 | 0.58 |
| **Pctl. 75** | 0.23 | 0.19 | 0.58 | 0.23 | 0.19 | 0.58 | 0.23 | 0.19 | 0.58 |
| **Max.** | 0.24 | 0.19 | 0.59 | 0.24 | 0.20 | 0.59 | 0.24 | 0.20 | 0.59 |

**Supplementary Table 2: Biopsy results for the simulations selected by ABC.** For all simulated levels of dispersal, the biopsy results closely matched the targets from the published data, supporting the model fit (Capalbo et al. 2021).

| Dispersal | Total aneuploid biopsies | Number of aneuploid biopsies originating from mosaic embryos |
|---|---|---|
| 0 | 581518 | 183313 (31.5%) |
| 0.5 | 581303 | 7891 (1.4%) |
| 1 | 583075 | 3975 (0.7%) |

**Supplementary Table 3: Proportion of aneuploid biopsies (i.e., biopsies where all 5 sampled cells were aneuploid) obtained from embryos that were actually mosaic.** All embryos used for biopsy were from the posterior predictive samples generated from the distribution of meiotic and mitotic error rates selected at each dispersal level.

| Embryo type | First biopsy | Second biopsy | Dispersal = 0 | Dispersal = 0.5 | Dispersal = 1 |
|---|---|---|---|---|---|
| Mosaic aneuploid | Euploid | Euploid | 108408 (47%) | 150107 (64.9%) | 156474 (68%) |
| Mosaic aneuploid | Euploid | Mosaic | 67793 (29.4%) | 80452 (34.8%) | 73288 (31.9%) |
| Mosaic aneuploid | Euploid | Aneuploid | 54450 (23.6%) | 702 (0.3%) | 49 (0%) |
| Mosaic aneuploid | Mosaic | Euploid | 70813 (37.7%) | 79995 (42.7%) | 77195 (41.3%) |
| Mosaic aneuploid | Mosaic | Mosaic | 59526 (31.7%) | 102154 (54.5%) | 106954 (57.2%) |
| Mosaic aneuploid | Mosaic | Aneuploid | 57492 (30.6%) | 5272 (2.8%) | 2804 (1.5%) |
| Mosaic aneuploid | Aneuploid | Euploid | 55148 (9.5%) | 759 (0.1%) | 72 (0%) |
| Mosaic aneuploid | Aneuploid | Mosaic | 57261 (9.8%) | 5410 (0.9%) | 2891 (0.5%) |
| Mosaic aneuploid | Aneuploid | Aneuploid | 70904 (12.2%) | 1722 (0.3%) | 1012 (0.2%) |
| Fully aneuploid | Aneuploid | Aneuploid | 398205 (68.5%) | 573412 (98.6%) | 579100 (99.3%) |
| Fully euploid | Euploid | Euploid | NA | 15 (0%) | 161 (0.1%) |

**Supplementary Table 4: First and second biopsy summaries of embryos constructed from the posterior distributions of meiotic and mitotic error rates inferred with ABC.** At each dispersal level, we also report the percentages of each combination of embryo type, first biopsy type, and second biopsy type out of all embryos of a given first biopsy type

| Data set | Maternal Age |
| --- | --- |
| Capalbo et al. (2021) | Mean: 38.06<br>Standard deviation: 3.65 |
| Clarke et al. (2023) | Median: 34.0<br>670 patients < 35<br>462 patients > 35 |
| Munné et al. (2017) | Mean: 35.8 |
| Rodrigo et al. (2020) | Mean: 38.6<br>Standard deviation: 6.2 |

**Supplementary Table 5: Maternal ages of PGT-A patients across input datasets.** Studies differed in reported measures of central tendency and variability.

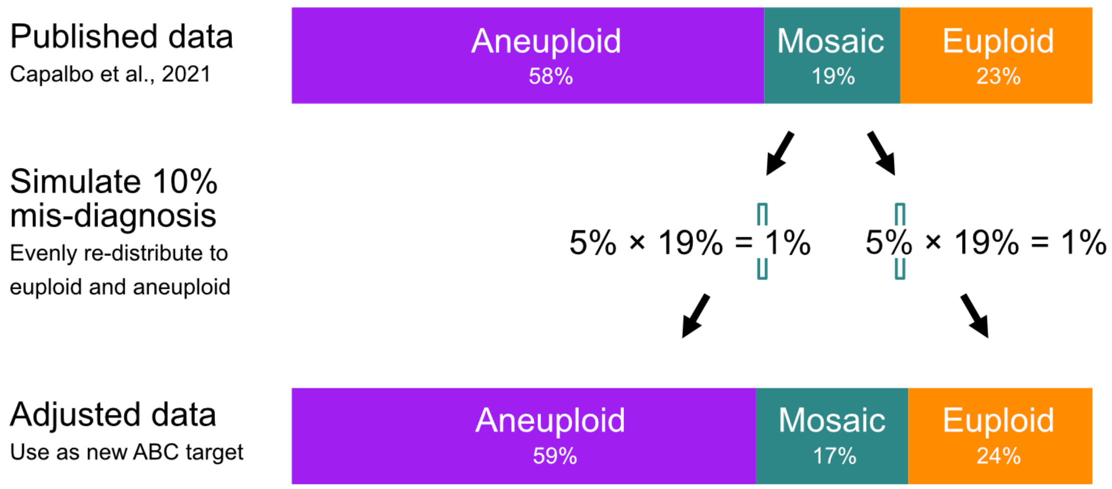


**Supplementary Figure 1: Schematic of approach for simulating misclassification of mosaic embryos.** We simulated varying levels of misclassification spanning from 0% to 100% in 10% increments. At each misclassification level, half of the mis-classified mosaic biopsies were assumed to be euploid and the other half aneuploid. These adjusted values were used as the new targets for ABC. The figure depicts an example of simulating a misclassification rate of 10% (Capalbo et al. 2021).
